# Fibrotic Extracellular Matrix Preferentially Induces a Partial Epithelial-Mesenchymal Transition Phenotype in a 3-D Agent Based Model of Fibrosis

**DOI:** 10.1101/2024.06.18.599477

**Authors:** Kristin P. Kim, Christopher A. Lemmon

## Abstract

One of the main drivers of fibrotic diseases is epithelial-mesenchymal transition (EMT): a transdifferentiation process in which cells undergo a phenotypic change from an epithelial state to a pro-migratory state. The cytokine transforming growth factor-*β*1 (TGF-*β*1) has been previously shown to regulate EMT. TGF-*β*1 binds to fibronectin (FN) fibrils, which are the primary extracellular matrix (ECM) component in renal fibrosis. We have previously demonstrated experimentally that inhibition of FN fibrillogenesis and/or TGF-*β*1 tethering to FN inhibits EMT. However, these studies have only been conducted on 2-D cell monolayers, and the role of TGF-*β*1-FN tethering in 3-D cellular environments is not clear. As such, we sought to develop a 3-D computational model of epithelial spheroids that captures both EMT signaling dynamics and TGF-*β*1-FN tethering dynamics. We have incorporated the bi-stable EMT switch model developed by Tian et al. (2013) into a 3D multicellular model, capturing both temporal and spatial TGF-*β*1 signaling dynamics. We show that the addition of TGF-*β*1 affects cell proliferation, EMT progression, and cell migration. We then incorporate TGF-*β*1-FN fibril tethering by locally reducing the TGF-*β*1 diffusion coefficient as a function of EMT to simulate the reduced movement of TGF-*β*1 when tethered to FN fibrils during fibrosis. We show that incorporation of TGF-*β*1 tethering to FN fibrils promotes a partial EMT state, independent of exogenous TGF-*β*1 concentration, indicating a mechanism by which fibrotic ECM can promote a partial EMT state.

**Author summary:** Epithelial-mesenchymal transition (EMT) is a key cellular process where epithelial cells transform and become mesenchymal. The EMT states are not binary, but instead exhibit a spectrum of partial states where epithelial cells express a combination of epithelial and mesenchymal markers, along with several markers distinct to the partial state. In diseases such as fibrosis and cancer, growing evidence supports the finding that diseased epithelial cells exist primarily in a partial EMT state. However, the mechanisms and signaling factors that drive this partial EMT state in fibrotic diseases is unclear. We use an agent-based model that looks at EMT progression in a population of cells embedded in an ECM environment with controllable fibrotic properties to provide a more systematic approach at studying spatial and temporal changes in the microenvironment that could drive EMT progression and maintain specific EMT phenotypes.

## Introduction

Epithelial-Mesenchymal transition (EMT) is a transdifferentiation process where epithelial cells shift to a mesenchymal-like phenotype [1–3]. Epithelial cells lose key phenotypic markers such as strong cell-cell adhesions, apicobasal polarity, and a cobblestone morphology, and acquire mesenchymal characteristics such as actin stress fibers, front-back polarity, and increased cell size [4]. EMT plays a major role in embryogenesis, tissue morphogenesis, and wound healing [5, 6]. In addition to the epithelial and mesenchymal states, studies have also identified an intermediate phenotype known as the partial state; this phenomena is also referred to as EMT plasticity [4, 7, 8]. In this partial EMT state, cells express a combination of epithelial and mesenchymal phenotypes and exhibit the ability to either revert to an epithelial state or proceed to the mesenchymal state in response to stimuli [5]. This partial state has been implicated in many EMT-associated diseases including cancer metastasis and fibrosis [1, 9–11]. Due to the reversibility of the partial EMT state, there has been increased interest in improving understanding of EMT plasticity and the mechanisms that drive it.

One of the most potent activators of EMT is the cytokine transforming growth factor-*β*1 (TGF-*β*1) [12, 13]. It is highly upregulated during metastasis as well as fibrosis in the kidney and liver [14, 15]. TGF-*β*1 drives upregulation of transcription factors Snail1 and Zeb1 [16,17]. Transcription factors Snail1 and Zeb1 repress expression of epithelial markers such as E-cadherin and promote expression of mesenchymal markers such as Vimentin and N-Cadherin [4, 16]. Snail has been shown to disrupt polarity of tubular epithelial cells during renal fibrosis, translocating from the nucleus to the plasma membrane, where it disrupts CRUMBS signaling [14]. Expression of Snail1 and Zeb1 are regulated via micro-RNAs (miRNA), with miRNA-34 and miRNA-200 widely identified as the main downregulators, respectively [16, 18]. TGF-*β*1 signaling increases expression of Snail1 and Zeb1 RNA, which in turn downregulates miRNA expression, subsequently leading to upregulation of transcription factor expression, ultimately driving upregulated mesenchymal marker expression and downregulated epithelial markers.

EMT is associated with alterations to the extracellular matrix (ECM); these ECM alterations in turn drive further EMT signaling (reviewed in [19]). Production of pro-fibrotic ECM proteins such as Collagen I and fibronectin are increased in EMT-associated diseases like fibrosis and cancer metastasis [14, 20]. TGF-*β*1 specifically drives expression and assembly of the ECM protein fibronectin (FN), which acts as a scaffold for cells during migration [21]. Soluble FN binds to integrins on the cell surface [20, 22]. Bound FN is then stretched in response to actomyosin forces transmitted via integrins that reveal cryptic FN-FN binding sites that promote fibril formation. Fibril formation also opens heparin binding domains that bind to over 40 different growth factors, including latent TGF-*β*1 [23, 24]. TGF-*β*1 binds to these domains on FN fibrils, colocalizing at the cell surface and promoting activation of TGF-*β*1 and subsequent TGF-*β*1 signaling [21, 25–27]. This creates a positive feedback loop of excessive ECM production and TGF-*β*1 signaling that drives fibrogenesis and promotes EMT progression.

Although *in vitro* experiments provide important physiological information on EMT progression, the time and cost of experiments makes it difficult to conduct long-term multivariate studies when performed in more complex culture models, such as 3D organoids or *in vivo* studies [28, 29]. There have been several computational models developed to study and map the TGF*β*1 signaling pathway, ranging from whole pathway signaling to reduced models that focus on key molecules that drive epithelial and mesenchymal marker expression [4, 16, 17, 30]. Several models have focused on the role of transcription factors Snail1 and Zeb1 and microRNA-34 (miR-34) and microRNA-200 (miR-200) in promoting EMT progression; these consist of a Snail/miR-34 negative feedback loop and a Zeb1/miR-200 negative feedback loop [16, 17]. Lu et al. modeled EMT as a ternary switch, where high levels of miRNA defined the epithelial state, high levels of Snail1 and Zeb1 represented the mesenchymal state, and an intermediate level expression of all 4 represented the partial state. The Tian model also focused on these 2 feedback loops via a bi-stable switch, where Snail1-miR-34 mediated the first transition to a partial state, and Zeb1-miR-200 mediated the second transition to a fully mesenchymal state. Several follow-up studies have built on these double negative feedback loops to improve predicted changes in cell state through increased complexity of TGF-*β*1 signaling or addition of other pathways that affect TGF-*β*1 signaling [31–33].

There have also been several models that investigated EMT using a multicellular approach to capture changes in cell-cell interactions that drive EMT. Bocci et al. accounted for cell-cell interactions by defining cells in a lattice, where neighboring cells influence their surroundings by secreting factors that can bind to neighbor cell receptors [33]; however, these simulations are constrained to 2-D models. The Cellular Potts model has been used to study spatial dynamics and the role of cell-cell junctional forces in tissue formation [34]. Recent work has also implemented the Cellular Potts model to study cell-cell and cell-matrix interactions that drive EMT [19]. Further implementation of this model also incorporated biochemical signaling [35]. However these models again focus on monolayers of cells, with no 3-D considerations. Furthermore, none of these models account for the effects of fibrotic ECM, which contains TGF-*β*1 binding sites on FN fibrils, which undoubtedly affects TGF-*β*1 signaling and diffusion.

In the current work, we address these shortcomings by incorporating the system of ordinary differential equations (ODEs) developed by Tian et al. [16] into a 3-D multicellular model that also simulates ECM assembly. Cells were initialized in a hollow-spheroid geometry, mimicking the cellular organization observed in *in vitro* spheroid experiments [36, 37], and FN assembly/ TGF-*β*1 tethering was modeled by locally altering the TGF-*β*1 diffusion coefficient depending on each cell’s EMT state: a reduction in local TGF-*β*1 diffusion coefficient simulates local ECM assembly and TGF-*β*1 tethering.

## Materials and methods

We developed an agent-based model that simulates the progression of EMT in a 3D multicellular geometry (Fig 1). The model integrates six (6) components to fully represent EMT progression in a 3D tissue: (1) initializing cells in a hollow spheroid geometry, (2) modeling intracellular TGF-*β*1 signaling, (3) classifying EMT state, (4) altering cell behavior as a function of EMT state, (5) changing local diffusion coefficients as a function of EMT state to simulate ECM remodeling, and (6) modeling extracellular TGF-*β*1 diffusion.

**Fig 1.**
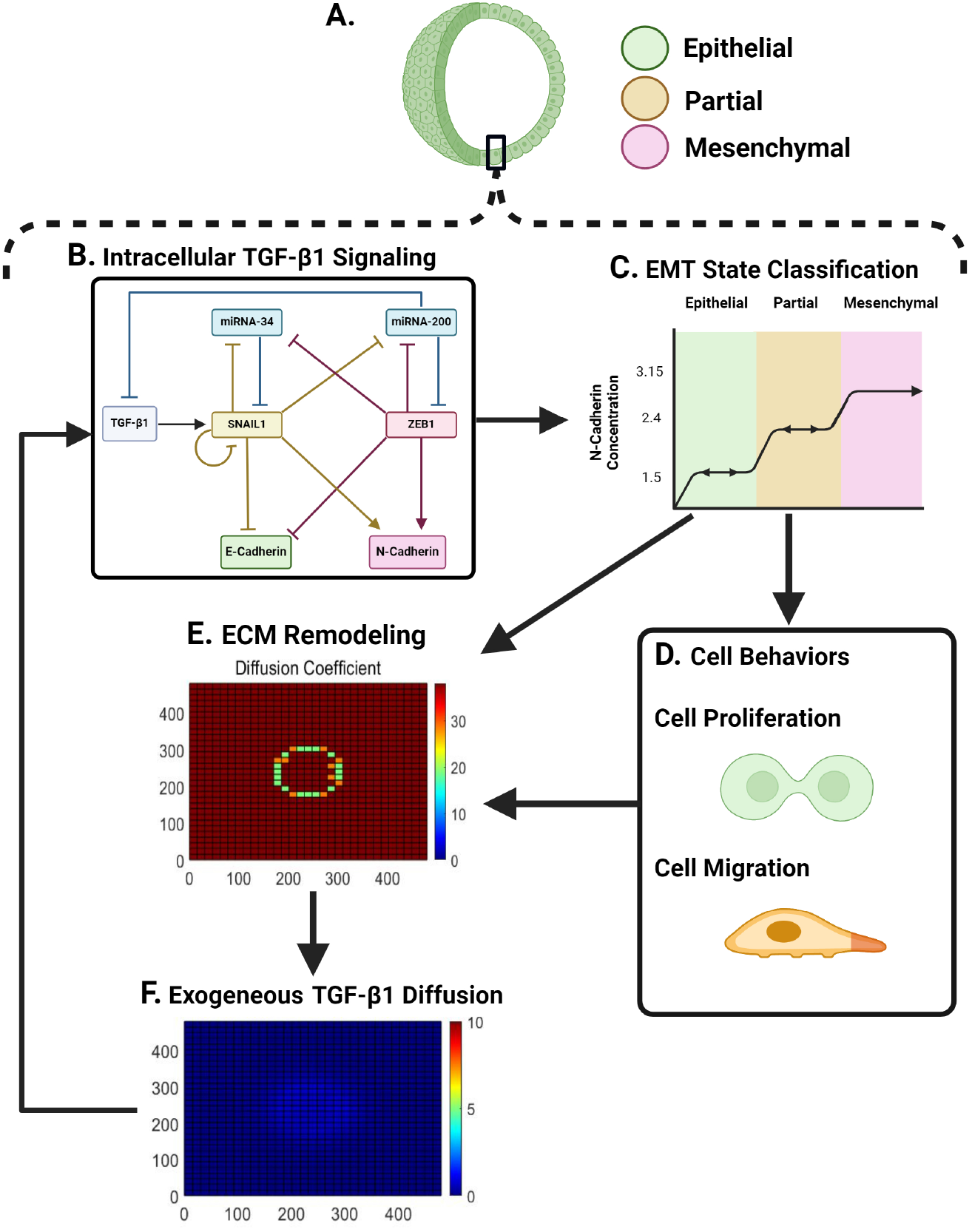
Model algorithm and organization. A: The model is initialized as a population of epithelial cells in a hollow spheroid geometry. B: Each cell has an integrated system of ordinary differential equations (ODEs) that define TGF-*β*1 dynamics within the cell. C: Outputs from the ODE system are used to classify EMT state (epithelial/partial/mesenchymal). D: Cell proliferation, migration, and TGF-*β*1 secretion are determined based on each cell’s EMT state. E: Local TGF-*β*1 diffusion coefficients are altered based on EMT state, with a lower diffusion coefficient simulating local ECM assembly. F: Diffusion coefficients are incorporated into a partial differential equation that simulates TGF-*β*1 diffusion in the system. Figure created using BioRender.com.

### Cell Agent-Based Model

We used an agent-based model to define the 3D multicellular spheroid, where each cell occupies a voxel with edge lengths of 15 µm, approximating the size of an average epithelial cell [38]. Cells can move from voxel to voxel, and cell location is tracked over time. We define an inner and outer radius to form a single-cell layered hollow spheroid, representative of cell geometries observed in *in vitro* epithelial spheroid studies [36, 39]. The total size of the model domain used is 32 x 32 x 32 voxels, correlating to a volume of 0.11 mm^3^. Initial investigations also looked at different tissue structures utilizing the same parameters shown above, which are discussed in the results.

### Intracellular TGF-*β*1 Signaling

For each cell, a system of ordinary differential equations was incorporated (Table S1). This ODE model simulates intracellular TGF-*β*1-induced signaling dynamics that drive EMT progression, where transcription factors Snail1 and Zeb1 serve as switches that promote epithelial transition to a partial state, and partial to a mesenchymal state, respectively. Initial concentrations and rate constants for the system of ODEs were defined based on previously published work [16] (Table S1, Table S2). The integrated bi-stable switch model has a constant, exogenous TGF-*β*1 concentration as an input. However, mesenchymal cells secrete *de novo* TGF-*β*1 [40, 41] into the extracellular space, which is a critical aspect of TGF-*β*1 dynamics that we sought to capture. As such, we assume immediate TGF-*β*1 secretion from cells and combine both endogenous and exogenous TGF-*β*1 into a single variable. As this differs from the original bi-stable switch model, we performed parameterization to estimate reaction kinetics that is discussed in the Supplemental Information.

### EMT Characterization

EMT progression in each cell was determined based on N-Cadherin concentration, which is an output from the intracellular TGF-*β*1 ODEs. Maximum N-Cadherin concentration is defined as 3.1515 µM (adapted from [35]). Cells were considered epithelial when N-Cadherin is *<* 1.5 µM, partial when N-Cadherin is between 1.5 µM and 2.4 µM, and mesenchymal when N-Cadherin is *≥* 2.4 µM. Previous work has shown that full EMT progression is an irreversible process [42], so cells in the mesenchymal state were constrained to remain mesenchymal. However, cells in a partial state are able to revert back to an epithelial phenotype [4, 42]; as such, we allowed cells in the partial state to return to the epithelial state if the N-Cadherin concentration of the inidividual cell dropped below 1.5 µM.

### Modeling Cell Proliferation and Migration

Rates of proliferation and migration were based on probability thresholds that depend on the cell state, where mesenchymal cells are more likely to migrate compared to epithelial cells (Table S3). Cell interactions were limited using a 3D Moore neighborhood, defined as a 3 x 3 x 3 window with the target cell set in the center position. Within this window, we defined the TGF-*β*1 concentrations and the position of any cells occupying neighboring voxels. The probability of cell migration is governed by a probability threshold dependent on the EMT state as previously done [35], where mesenchymal cells were given higher probability of migration compared to epithelial (Table S4). We further set a rule that promoted cell migration to the neighboring unoccupied voxel with the highest TGF-*β*1 concentration.

To model cell proliferation, cells were set to have a baseline rate of one division per every 24 hours. Once this time point was reached, we again used a probability threshold to drive proliferation based on the EMT state, where cells in the epithelial state are more likely to divide relative to those in the mesenchymal state (Table S4) The position of the new daughter cell was decided based on free space present in the Moore neighborhood, and the new daughter cell acquired the same parameters and EMT state as the parent cell.

### ECM Remodeling

One of the interesting dynamics in TGF-*β*1 signaling is the positive feedback loop developed between TGF-*β*1 and FN [27]. Increased TGF-*β*1 activation leads to increased synthesis of FN, which then feeds back and promotes localization of TGF-*β*1 to the assembled ECM [21]. FN synthesis and fibril formation were modeled through local changes in the TGF-*β*1 diffusion coefficient, where each voxel is assigned a diffusion coefficient value that is tracked through each time step. The initial diffusion coefficient value of each position was set to 8550 µm^2^/hr. Given that each voxel is 15x15 *µm*^2^, a baseline diffusion coefficient of 38 pixels^2^/hr was used in simulations. This baseline diffusion coefficient is consistent with experimental values of growth factor diffusion in collagen gels [43]. Physiologically, increased FN assembly and TGF-*β*1 tethering to FN fibrils should decrease TGF-*β*1 diffusion through the ECM [23,44,45]. To simulate changes in ECM remodeling, we considered the dependence of cell state on ECM assembly. As cells undergo EMT, we expect increased FN assembly, therefore we expect decreased TGF-*β*1 diffusion. To implement this in the code, we use the following equation:

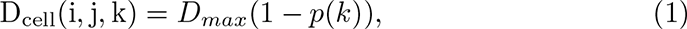

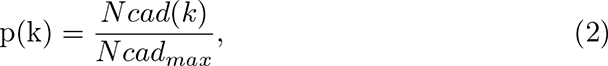

where the diffusion coefficient decreases based on the ratio of N-Cadherin, ensuring gradual decrease in diffusion coefficient value and subsequent representation of ECM remodeling as cells undergo EMT.

### Extracellular TGF-*β*1 Diffusion

Mimicking standard *in vitro* protocols, an initial concentration of exogenous TGF-*β*1 was applied to cells in the model. Initial TGF*β*1 concentration was assumed to be equally distributed in each voxel. For each time step, TGF-*β*1 diffusion is governed by a diffusion equation [46]:

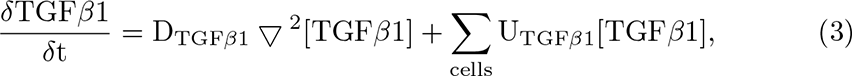

where [TGF-*β*1] is the concentration of TGF-*β*1, D_TGF-*β*1_ is the diffusion coefficient, and U [TGF-*β*1] is the cell contribution to the TGF-*β*1 concentration (cells are able to produce or degrade the cytokine based on the TGF-*β*1 conservation equation within the intracellular system of ODEs). To prevent cell migration out of the model domain, a boundary condition of [TGF-*β*1] = 0 at boundaries was applied.

### Computational Analysis

All simulations were performed in MATLAB. For each experiment, we averaged the output values for 20 simulations per condition. Graphical data is expressed as means *±* SEM. Statistical differences between conditions were assessed using either 1-way or 2-way ANOVA in GraphPad Prism 10, as noted on individual figures. P-values were used to determine statistical significance and are shown in figure legends, where significance was defined using *α* = 0.05.

To facilitate open use of the model and for easier access for future research, we translated the original code into a GUIDE using the MATLAB App Designer. Further details and examples of the application outputs are shown in Figure S4.

## Results

### Effect of tissue architecture on EMT progression

Initial simulations investigated whether changes in EMT progression were in agreement with previous published *in vitro* work and computational models. Simulations were run using one of three tissue architectures: i) a hollow spheroid; ii) a hollow cylinder; and iii) a curved cylinder (Fig 2A). For all three geometries, epithelial cells were initially organized with an outer radius of 4.5 voxels and an inner radius of 4.0 voxels, thus ensuring single cell thickness. Creation of the curved tube morphology was done by adapting a previously published code (TubePlot) [47].

**Fig 2.**
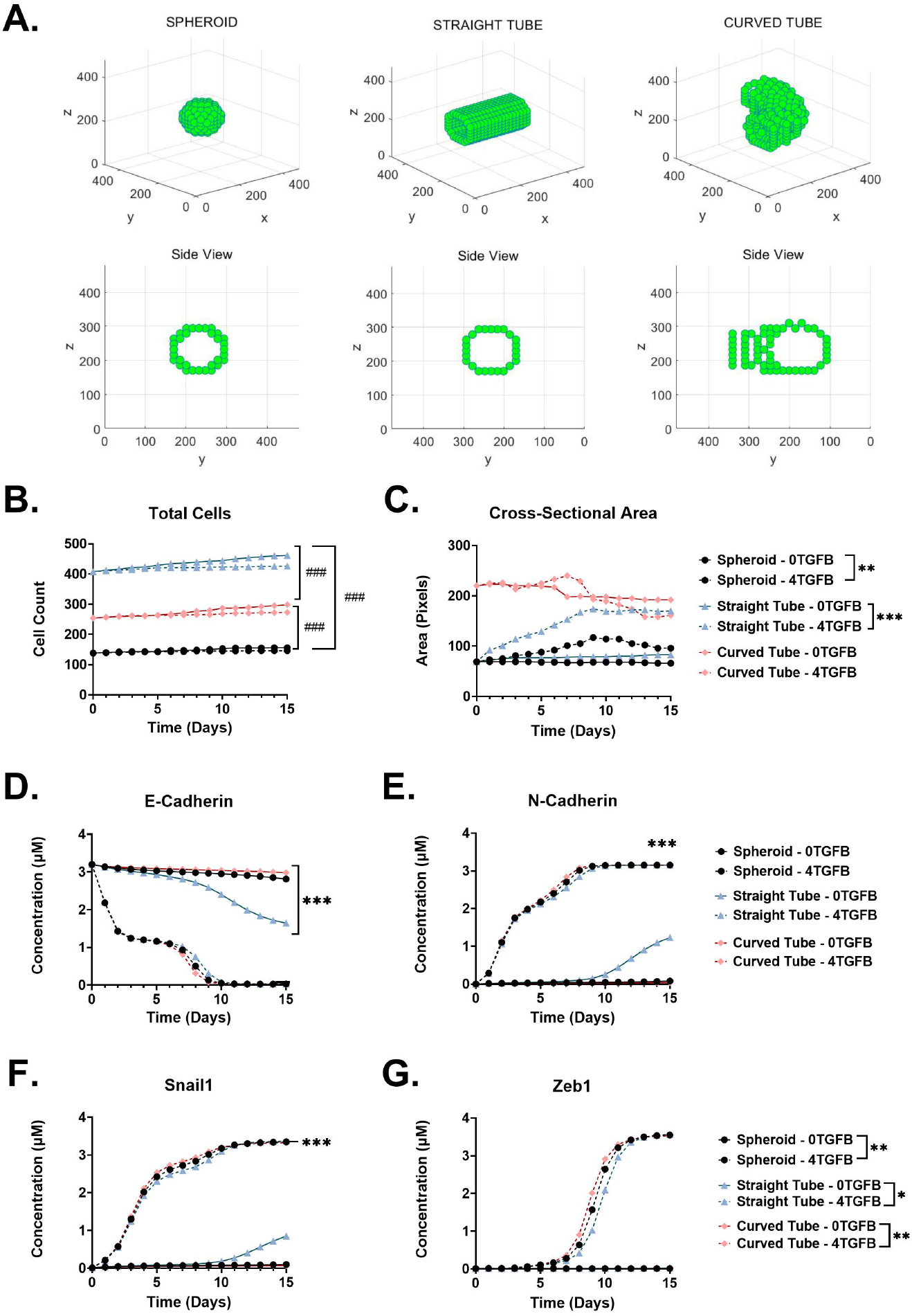
Simulating EMT in multiple tissue architectures. As a first investigation, the model was used to simulate EMT in 3 distinct tissue architectures (spheroid, straight tube, or curved tube). A) Representative images of the different tissue architectures. B) Total cell count as a function of time and tissue architecture when exposed to 0 or 4 µM exogenous TGF-*β*1. C) Cross-sectional area of the tissue architecture as a function of time and tissue architecture when exposed to 0 or 4 µM exogenous TGF-*β*1. D-G) Average concentrations of EMT markers (D) E-Cadherin, (E) N-Cadherin, (F) Snail, and (G) Zeb1. Statistics were analzyed using a 1-way ANOVA and post hoc Tukey multiple comparison test (*^∗^p ≤* 0.033, *^∗∗^p ≤* 0.002, *^∗∗∗^p ≤* 0.001; *^∗^*: Comparisons between TGF-*β*1 conditions; ^#^: Comparisons between morphology).

Simulations were run with cell locations initialized in each of the three tissue architectures in the absence or presence of 4 µM exogenous TGF-*β*1. Effects of tissue architecture are shown in Fig. 2B-G. Significant differences in total cell count were observed between the 3 architectures ((Fig 2B), which was expected given the differences in cell initialization, where the tubule-like structures require a higher initial cell count compared to the hollow spheroid. Changes in cross-sectional area also exhibited similar trends, where larger cross-sectional areas were observed in the tubular structures versus spheroids without any added TGF-*β*1. EMT progression was similar at early time points between all three tissue architectures (Fig 2C-F), with cells remaining epithelial in the absence of TGF-*β*1 and progressing to a mesenchymal state in the presence of TGF-*β*1. Differences in EMT state were observed at later time points, suggesting that tissue architecture may play a role in driving local EMT during longer time points regardless of TGF*β*1 stimuli. Given the prevalence of hollow spheroids as an *in vitro* experimental tool to investigate EMT in both development and disease states [39], we chose to focus on the spheroid morphology for the remainder of the current work, investigating cellular responses up to Day 10, where cells are expected to reach full mesenchymal-like state in response to 4 µM TGF-*β*1 stimuli. Results from simulations using tubular architectures are shown in supplementary materials (Figure S3, Figure S4).

### Effect of TGF-*β*1 concentration on EMT progression and tissue morphology

We next investigated the effects of TGF-*β*1 concentration on EMT progression in spheroids in the absence of a fibrotic ECM. Representative simulation outputs are shown in Fig. 3A, with cell locations, EMT state, TGF-*β*1 concentration, and TGF-*β*1 diffusion coefficient shown at the last time step and at a cross-sectional view through the centroid of the computational space. Representative simulation outcomes are shown for the entire computational time in Supplemental Movie S1. In the absence of exogenous TGF-*β*1, cells maintained a population-wide epithelial state (green), characterized by low migration and high rates of proliferation. Graphical representations also show low TGF*β*1 production and minimal changes to the surrounding D_TGF-*β*1_ values, suggesting low ECM assembly. Cells in spheroids exposed to 2 µM exogenous TGF-*β*1 initiated the transition to a partial EMT state (yellow), still with low TGF-*β*1 production, but increased ECM assembly. At 4 µM TGF-*β*1, cells transitioned to a full mesenchymal phenotype (red), with a significant increase in TGF*β*1 production and ECM remodeling (Fig 3A).

**Fig 3.**
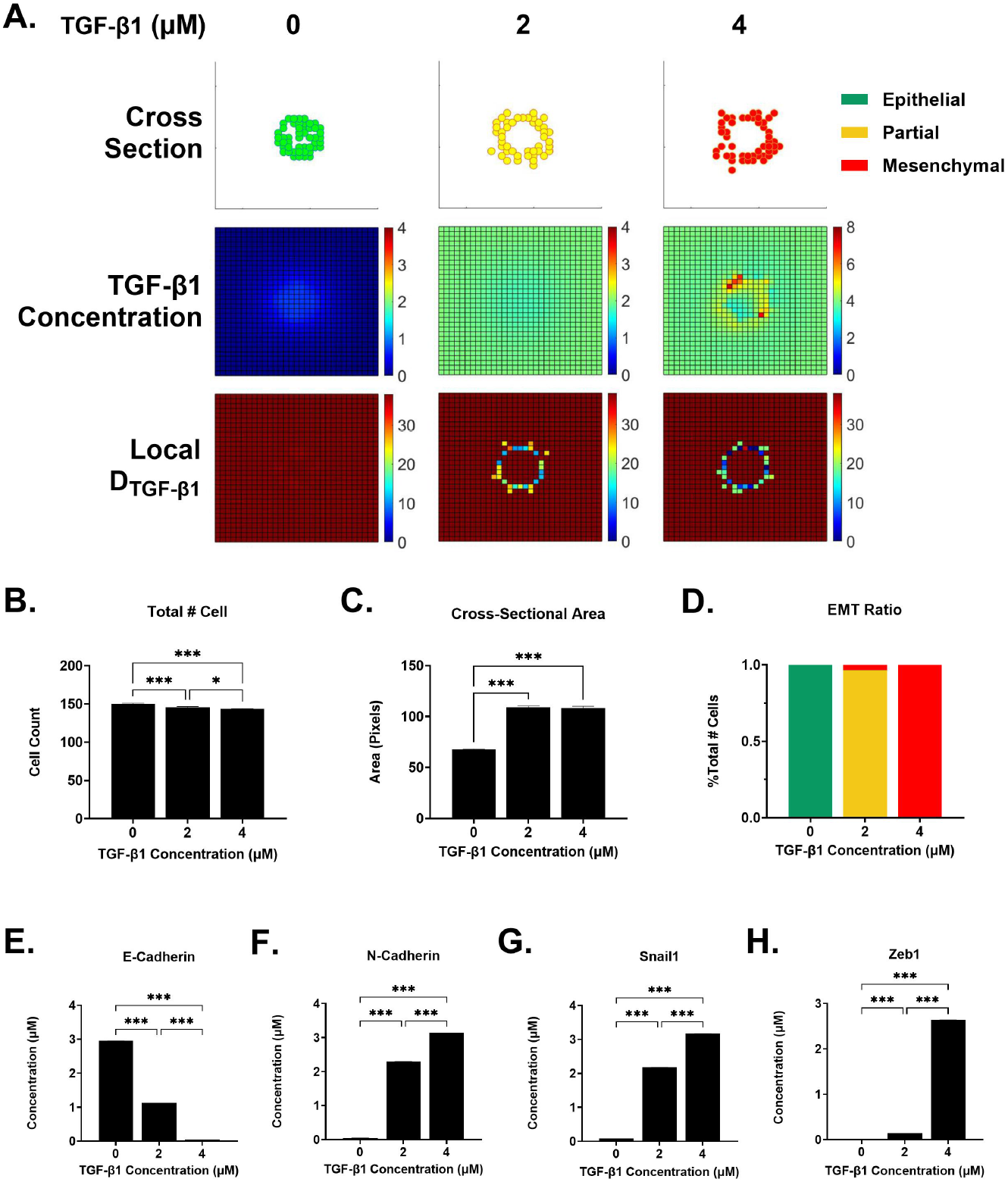
Exogenous TGF-*β*1 drives EMT progression. Cells were exposed to 0, 2, or 4 µM TGF-*β*1 A) Representative images of spheroids, TGF-*β*1 concentration, and D_TGF-*β*1_. B) Total cell count as a function of exogenous TGF-*β*1 concentration. C) Spheroid cross-sectional area as a function of exogenous TGF-*β*1 concentration. D) Fraction of cells in each EMT state (epithelial, partial, mesenchymal) as a function of exogenous TGF-*β*1 concentration. E-H) Changes in EMT markers as a function of exogenous TGF-*β*1 concentration: E) E-Cadherin, F) N-Cadherin, G) Snail, H) Zeb1. N = 20 for each condition. Statistics were analyzed using a 1-way ANOVA and post hoc Tukey multiple comparison test (*^∗^p ≤* 0.033, *^∗∗^p ≤* 0.002, *^∗∗∗^p ≤* 0.001).

Changes in spheroid morphology (Fig 3B,C), EMT state (Fig 3D)and EMT marker concentrations (Fig 3E-H) were averaged for all simulations. Without TGF-*β*1 stimuli, cells maintained their initial spheroid size and exhibited increased proliferation. As exogenous TGF-*β*1 increased, there was a significant decrease in cell proliferation, as observed by decreases in total cell count (Fig. 3B), and a significant increase in cell migration. Together, this drove an increase in the size of the lumen as cells migrated outwards from the initial spheroid shape (Fig. 3C).

Figure 3E-H shows the average concentration of EMT markers as a function of TGF-*β*1 concentration. Results were consistent with those observed in the original bi-stable switch model [16]. E-Cadherin decreased with increasing TGF-*β*1 concentration, while N-Cadherin increased. Snail exhibited a significant increase at 2 µM TGF-*β*1, consistent with a predominantly partial EMT state and remained elevated at 4 µM TGF-*β*1. Zeb1 exhibited a slight increase at 2 µM TGF-*β*1, and then a substantial increase at 4 µM TGF-*β*1, indicating a transition to the full mesenchymal state at the highest TGF-*β*1 concentration.

### Increased exogenous TGF-*β*1 leads to localized pools of TGF-*β*1 around the spheroid, corresponding to lower D_TGF-*β*1_ values

Representative changes in TGF-*β*1 concentration and TGF-*β*1 diffusion coefficient are shown in Fig 3A. In the absence of exogenous TGF-*β*1, the only TGF-*β*1 in the system is cell-produced TGF-*β*1, and as such is concentrated within the spheroid. At 2 µM exogenous TGF-*β*1, there is an observable consumption of exogenous TGF-*β*1, leading to a reduced concentration within the spheroid relative to the bulk concentration. Further increasing exogenous TGF-*β*1 to 4 µM led to increased cell production of TGF-*β*1 that began to localize at the spheroid periphery. As expected, these pools of high TGF-*β*1 values correlate to areas of low TGF-*β*1 diffusion coefficient values.

The local diffusion coefficient is defined based on the N-Cadherin concentration of each individual cell. Representative changes in the local TGF-*β*1 diffusion coefficient can be observed in Fig 3A. At 0 µM concentrations of exogenous TGF-*β*1, there is no observable decrease in *D_T GF β_*_1_, suggesting no ECM assembly. As exogenous TGF-*β*1 concentration increases, a decrease in *D_T GF β_*_1_ is observed around the spheroid periphery, consistent with previous findings that indicate a correlation between exogenous TGF-*β*1 and increased TGF-*β*1-FN tethering [27, 48]. This decrease in *D_T GF β_*_1_ also correlates spatiotemporally with EMT state, with a larger reduction in *D_T GF β_*_1_ correlating with the location of cells in a more mesenchymal state (Fig 3A).

### A fibrotic ECM induces the partial EMT state in the absence of exogenous TGF-*β*1

Simulations to this point were conducted with an initial high TGF-*β*1 diffusion coefficient, which models a non-fibrotic ECM. We next investigated the effects of the initial ECM environment; that is, if the spheroid is embedded in a fibrotic ECM, are there significant differences in the model outcomes? A fibrotic environment by definition has increased ECM assembly and deposition [49,50], with a particularly notable increase in FN fibril assembly [21, 51], which has been shown to tether TGF-*β*1 [44, 52]. Fibrosis is modeled in simulations by decreasing the initial diffusion coefficient of TGF-*β*1 in the system (D_TGF-*β*1_), resulting in reduced local diffusion of TGF-*β*1, as would be observed if TGF-*β*1 is tethered to FN. To simulate this, D_TGF-*β*1_ of the extracellular voxels was set to 38, 19, 4, or 0 pixels^2^/hr, without any exogenous TGF-*β*1. Representative simulation outcomes are shown at the final computational time point for a cross-sectional view through the computational space in Fig4A. As D_TGF-*β*1_ decreases, an increase in EMT is observed, where the partial EMT state occurs at lower D_TGF-*β*1_ values (Fig4A). To further examine EMT state in this model, the ratio of cells in each state (epithelial/partial/mesenchymal) is shown as a function of time (Fig. 4B), and the previously examined EMT markers are also shown as a function of time (Fig. 4C). At lower values of D_TGF-*β*1_, cells remain predominantly epithelial until 7-10 days of exposure to the fibrotic environment, with a sharper transition to a partial EMT state observed when D_TGF-*β*1_ is equal to 0. EMT markers increase with time as D_TGF-*β*1_ decreases, with markers reaching a steady state when D_TGF-*β*1_) is equal to 0.

**Fig 4.**
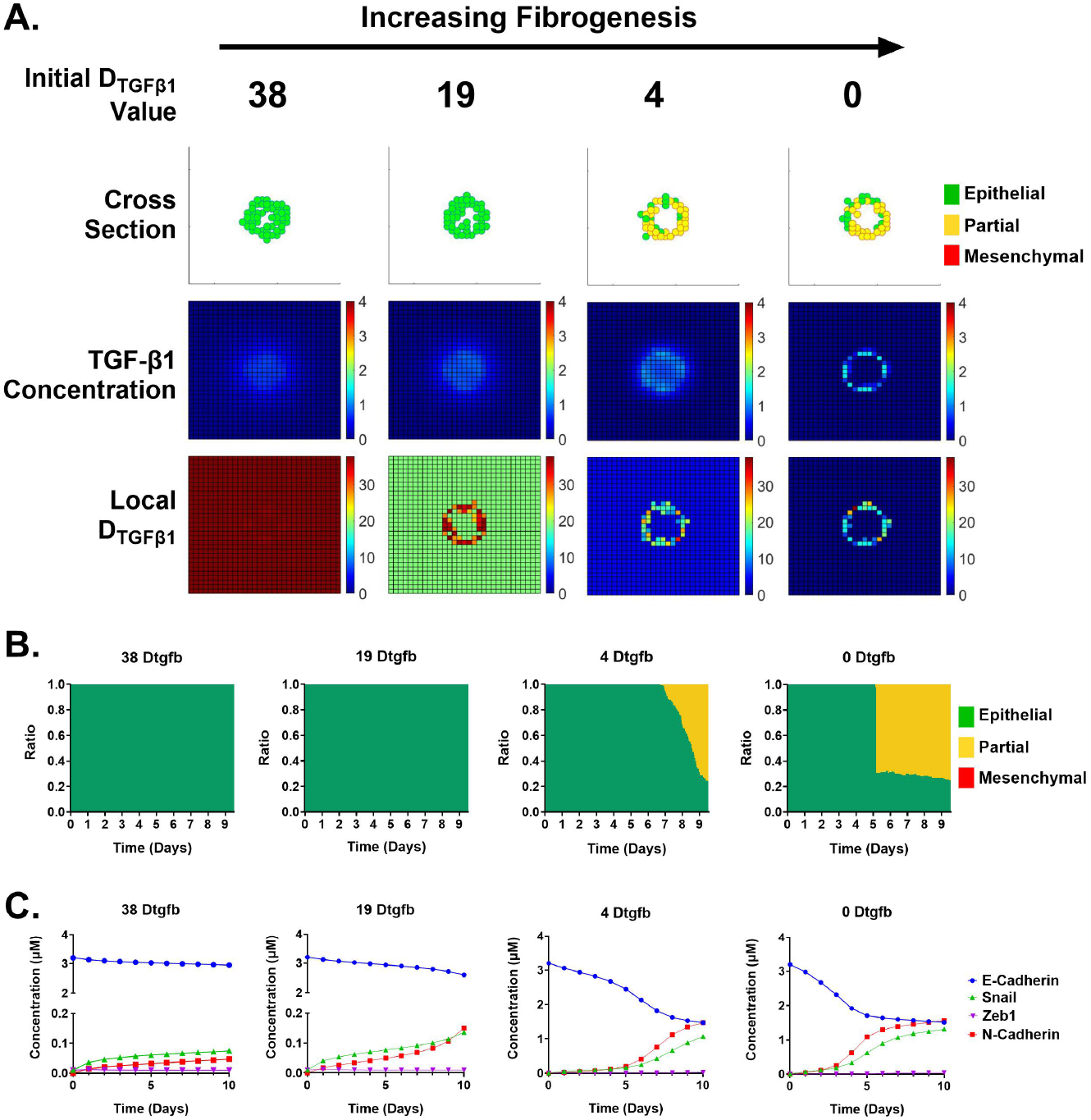
Effects of increased fibrotic ECM in the absence of exogeneous TGF-*β*1. Changes in the fibrotic state of the ECM were modeled by decreasing D_TGF-*β*1_. Simulations were conducted using D_TGF-*β*1_ values of 38, 19, 4 or 0 pixels^2^/hr. A) Representative simulation outputs for cellular organization, TGF-*β*1 concentration, and D_TGF-*β*1_. B) Fraction of cells in each EMT state as a function of time and D_TGF-*β*1_. C) Changes in EMT markers as a function of time and D_TGF-*β*1_.

### A fibrotic ECM promotes the partial EMT state in the presence of exogenous TGF-*β*1

Given the results that a fibrotic ECM promotes the partial state in the absence of exogenous TGF-*β*1, we next investigated the effects of a fibrotic ECM on EMT in the presence of exogenous TGF-*β*1. An initial concentration of 4 µM TGF-*β*1 was applied to the system and again the initial D_TGF-*β*1_ was set to 38, 19, 4 or 0 pixels^2^/hr for all voxels outside of the spheroid. Representative simulation outputs are shown at the final computational time point and for a cross-sectional view through the centroid of the computational space in Fig5. At high D_TGF-*β*1_ values, cells transition to a fully mesenchymal state, consistent with earlier simulations. However, at lower initial D_TGF-*β*1_ values, we surprisingly see a heterogeneous populations of cells in the mesenchymal and partial states. EMT state (Fig. 5B) and average EMT marker values (Fig. 5C) are again calculated as a function of time. These indicate that at higher D_TGF-*β*1_ values, cells proceed through two state changes, from E to P and P to M. However, at lower D_TGF-*β*1_ values, the transition to mesenchymal is delayed and is eventually absent. EMT marker expression shows a near steady-state scenario in which the partial state is predominant.

**Fig 5.**
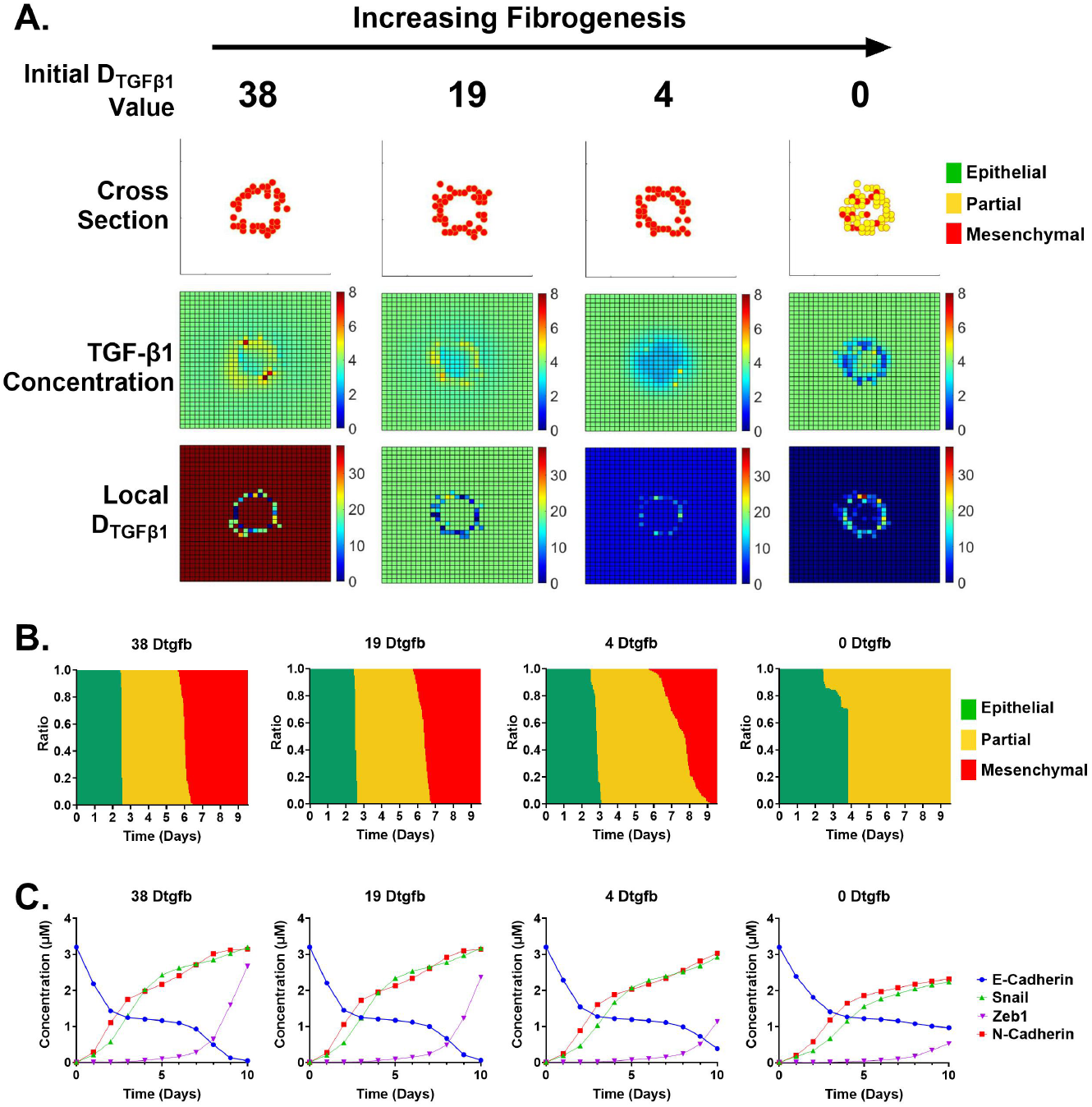
Effects of increased fibrotic ECM in the presence of 4 µM exogeneous TGF-*β*1. Changes in the fibrotic state of the ECM were modeled by decreasing D_TGF-*β*1_. Simulations were conducted using D_TGF-*β*1_ values of 38, 19, 4 or 0 pixels^2^/hr. A) Representative simulation outputs for cellular organization, TGF-*β*1 concentration, and D_TGF-*β*1_. B) Fraction of cells in each EMT state as a function of time and D_TGF-*β*1_. C) Changes in EMT markers as a function of time and D_TGF-*β*1_.

### Combinatorial effects of fibrotic ECM and exogenous TGF-*β*1

Results so far suggest that a fibrotic ECM promotes the partial EMT state in both the absence of exogenous TGF-*β*1 and presence of 4 µM exogenous TGF-*β*1. To further characterize these interactions, simulations were run in which we varied both exogenous TGF-*β*1 (0, 2 or 4 µM) and D_TGF-*β*1_ (38, 19, 4 or 0 pixels^2^/hr). The presence of a fibrotic ECM decreased cell count in the absence of exogenous TGF-*β*1, but had no significant effect in the presence of 2 or 4 µM exogenous TGF-*β*1 (Fig. 7). Spheroid cross-sectional area increased with increasing exogenous TGF-*β*1 (Fig. 6B); this increase was amplified by increasing fibrotic ECM in the absence of exogenous TGF-*β*1 but was inhibited by increasing fibrotic ECM in the presence of exogenous TGF-*β*1. EMT state (Fig. 6C) shows a clear transition from epithelial to partial to mesenchymal states in the presence of 0, 2 and 4 µM TGF-*β*1 in a non-fibrotic ECM, with a transition to a partial EMT state predominance at all concentrations of exogenous TGF-*β*1 with increasingly fibrotic ECM. This is supported by levels of the EMT markers (Fig. 6D-H). E-Cadherin significantly decreased with increasing fibrotic ECM in the absence of exogenous TGF-*β*1, but increased in the presence of exogenous TGF-*β*1; N-Cadherin and Snail1 showed inverse trends to E-Cadherin; and Zeb1, indicative of a transition to a fully mesenchymal state, was significantly inhibited by decreased D_TGF-*β*1_ at high exogenous TGF-*β*1.

**Fig 6.**
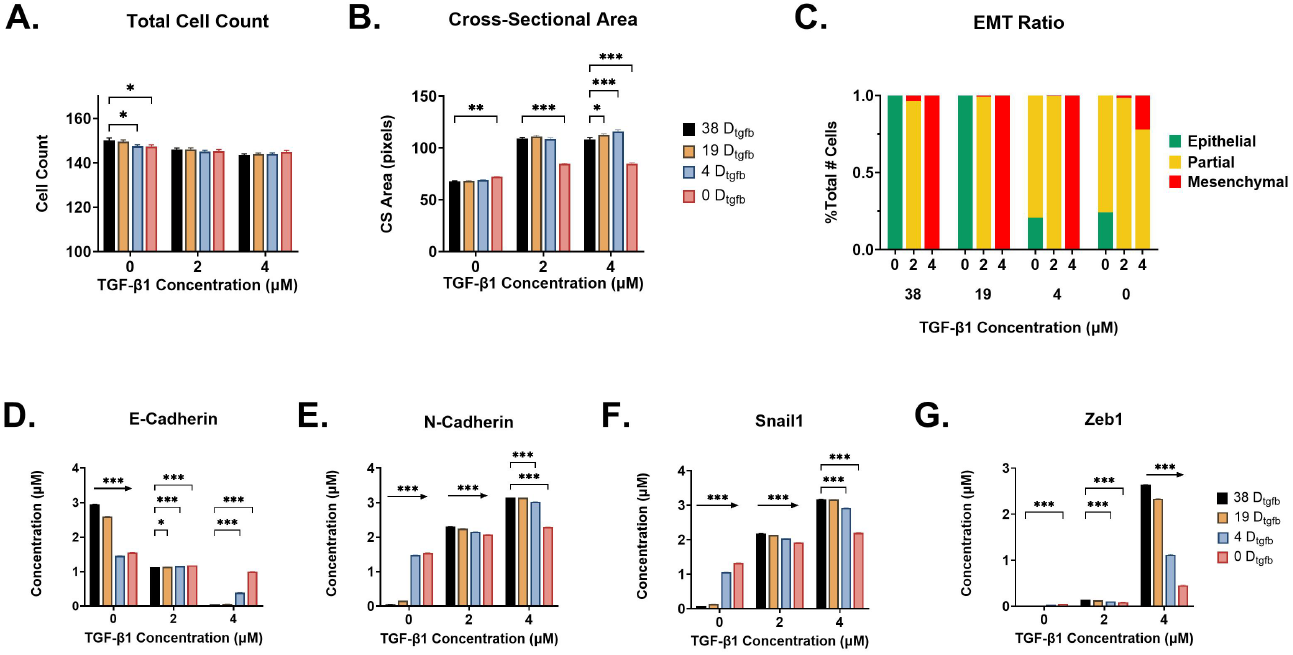
Combinatorial effects of exogenous TGF-*β*1 and fibrotic ECM. Cells were exposed to 0, 2 or 4 µM TGF-*β*1 and D_TGF-*β*1_ values of 38, 19, 4, or 0 pixels^2^/hr. A) Total cell count as a function of exogenous TGF-*β*1 and D_TGF-*β*1_. B) Spheroid cross-sectional area as a function of exogenous TGF-*β*1 and D_TGF-*β*1_. C) Fraction of cells in each EMT state as a function of exogenous TGF-*β*1 and D_TGF-*β*1_. D-G) Changes in EMT markers as a function of exogenous TGF-*β*1 and D_TGF-*β*1_: D) E-Cadherin; E) N-Cadherin; F) Snail1, and G) Zeb1. N = 20 simulations per condition. Statistics were run using a 2-way ANOVA and post hoc Tukey multiple comparison test (*^∗^p ≤* 0.033, *^∗∗^p ≤* 0.002, *^∗∗∗^p ≤* 0.001).

**Fig 7.**
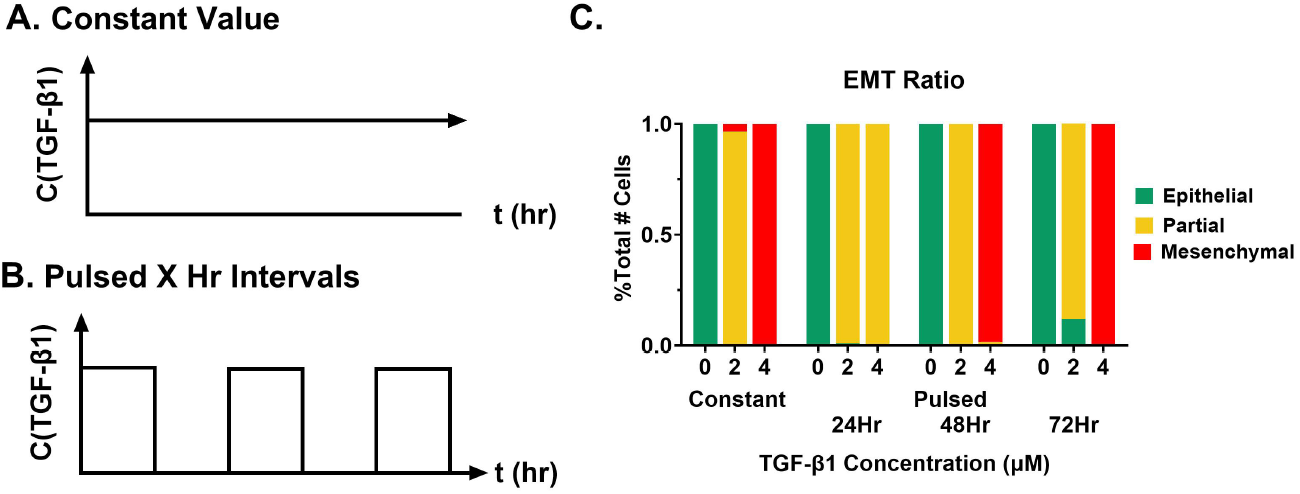
Modeling TGF-*β*1 exposure. A,B) Graphical representation of TGF-*β*1 exposure for A) constant and B) pulsed TGF-*β*1 exposure. C) Fraction of cells in each EMT state as a function of exogenous TGF-*β*1 concentration and pulse time. N = 20 simulations per condition).

### Effects of pulsed exogenous TGF-*β*1

All simulations to this point have been conducted in the presence of a constant initial concentration of exogenous TGF-*β*1. A frequent criticism of *in vitro* experimental work in EMT is that cells are exposed to constant TGF-*β*1, which may be inconsistent with oscillatory or pulsatile TGF-*β*1 exposure *in vivo*. Our model provides a platform to investigate the effects of pulsed versus constant TGF-*β*1 on EMT progression. Simulations were run in which TGF-*β*1 was either constant (Fig. 7A) or pulsed (Fig. 7B) with a pulse time of 24, 48, or 72 hours. Concentrations of 0, 2 and 4 µM TGF-*β*1 were used. Results suggest that pulsing at a 24 hr interval promotes the partial EMT state (Fig. 7C), while pulsing at longer intervals results in EMT states similar to constant TGF-*β*1 exposure. Representative simulation outputs are shown for the final time step through a cross-sectional view at the centroid of the computational space for both 2 µM TGF-*β*1 (Fig. 8A) and 4 µM TGF-*β*1 (Fig. 8B). These results indicate that despite similar changes in D_TGF-*β*1_, there is a marked predominance of cells in the partial EMT state for the 24 hour pulsed condition. This difference is particularly pronounced at 4 µM TGF-*β*1 (Fig. 8B).

**Fig 8.**
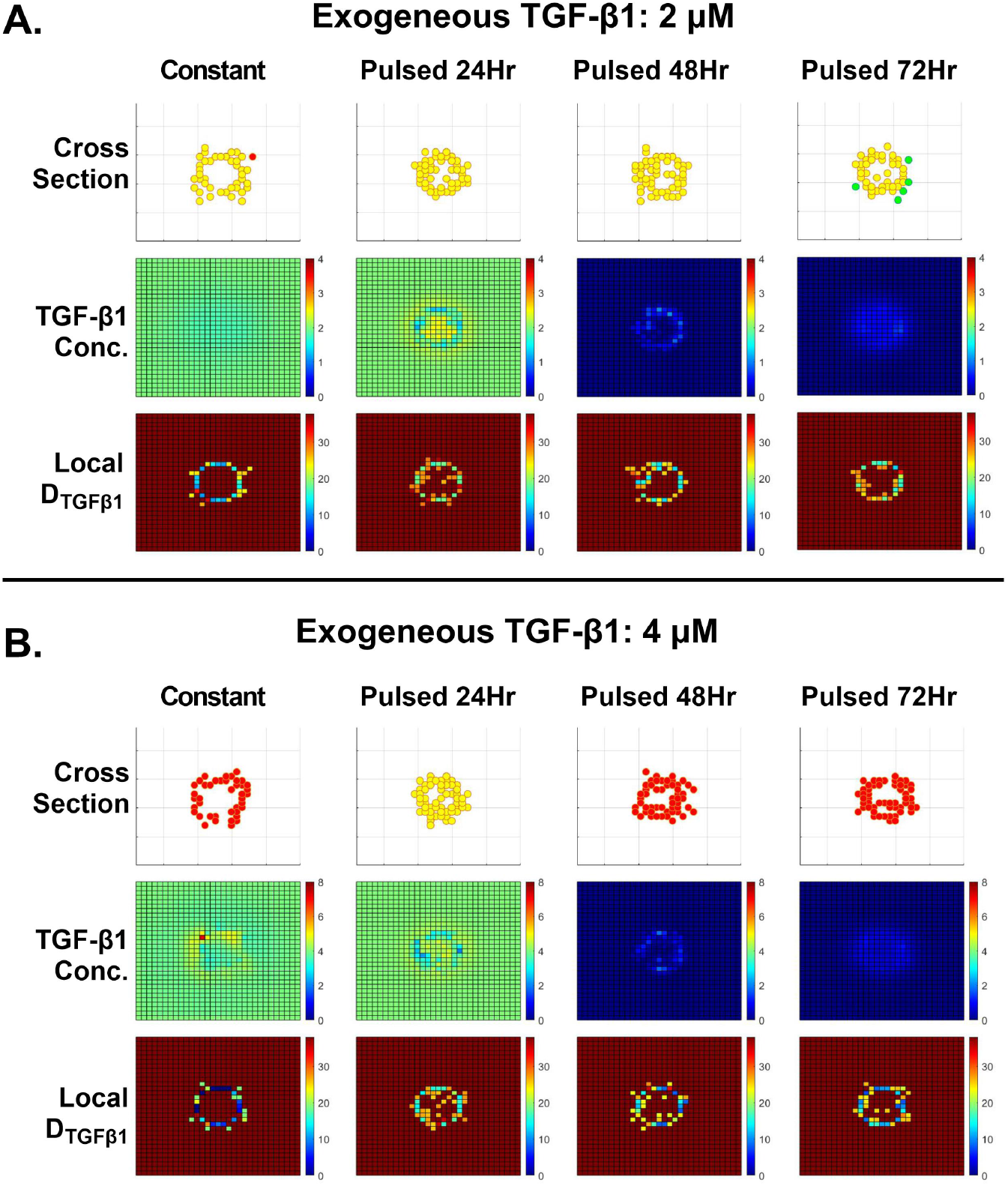
Effects of exogenous TGF-*β*1 concentration and pulsing. A) Representative outputs of cellular architecture, TGF-*β*1 concentration, and D_TGF-*β*1_ at 2 µM exogenous TGF-*β*1. B) Representative outputs of cellular architecture, TGF-*β*1 concentration, and D_TGF-*β*1_ at 4 µM exogenous TGF-*β*1.

We next examined effects of pulsatile versus constant TGF-*β*1 exposure on cell count (Fig. 9A), spheroid cross-sectional area (Fig. 9B), and EMT markers (Fig. 9C-F). The rate of pulsed TGF-*β*1 only affected cell count at 4 µM TGF-*β*1, where an increase in cell number with increasing pulse duration is observed. Pulsatile exposure decreased spheroid cross-sectional area at 2 and 4 µM TGF-*β*1. Effects of pulsatility on EMT markers showed a preference for the partial EMT state, which is particularly noted by the decreased Zeb1 expression (Fig. 9F). This suggests that pulsatile exogenous TGF-*β*1 results in partial EMT, possibly due to reversion of cells in a partial state back to an epithelial state.

**Fig 9.**
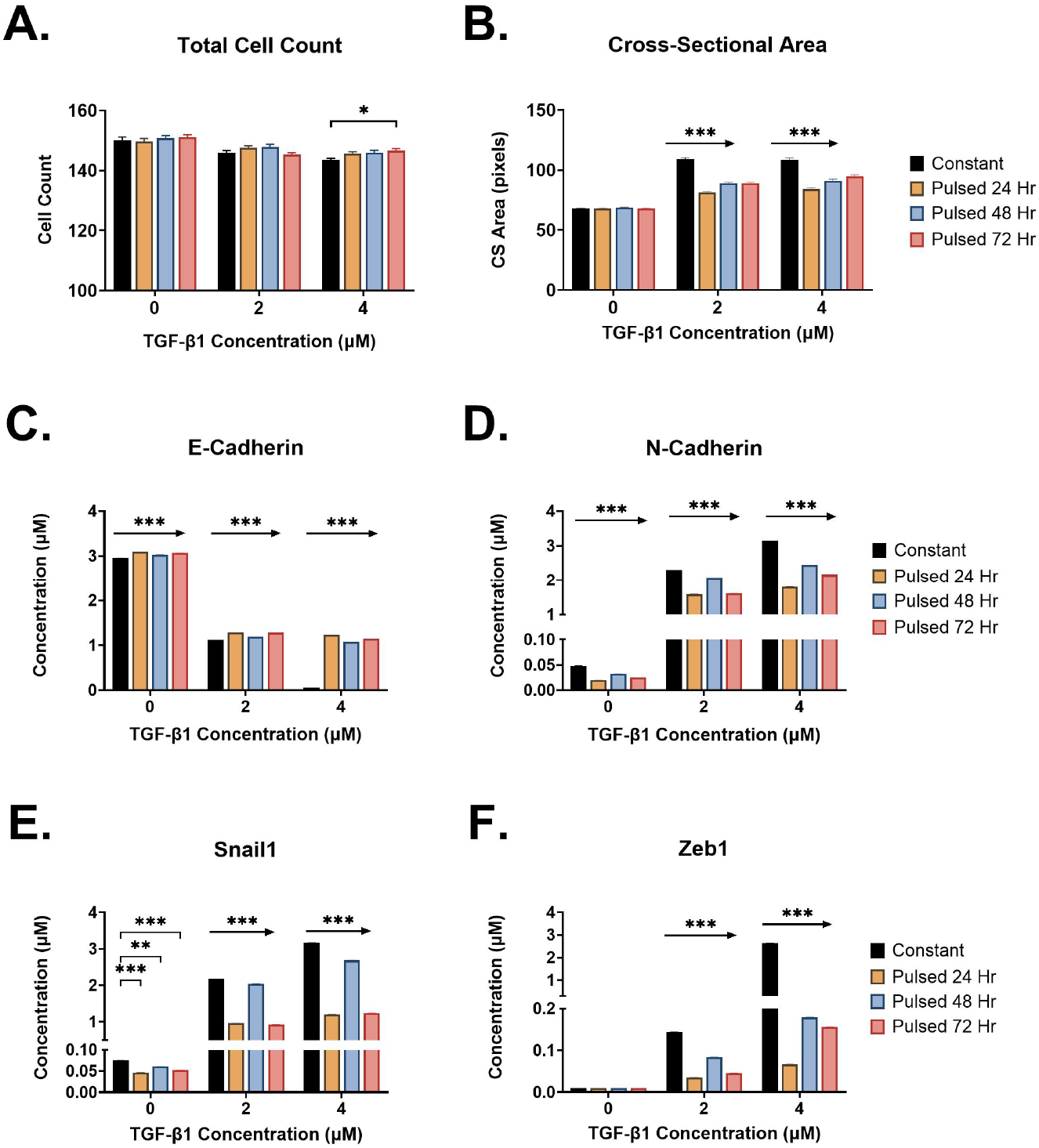
Effects of exogenous TGF-*β*1 concentration and pulsing. A) Total cell count as a function of exogenous TGF-*β*1 concentration and pulse rate. B) Spheroid cross-sectional area as a function of exogenous TGF-*β*1 concentration and pulse rate. C-F) EMT markers as a function of exogenous TGF-*β*1 concentration and pulse rate: C) E-Cadherin, D) N-Cadherin, E) Snail1, and F) Zeb1. N = 20 simulations per condition). Statistics were analyzed using a 1-way ANOVA and post hoc Tukey multiple comparison test (*^∗^p ≤* 0.033, *^∗∗^p ≤* 0.002, *^∗∗∗^p ≤* 0.001).

## Discussion

In this study, we developed a stochastic agent-based model of cells in a 3D tissue that can predict changes in EMT state in response to exogenous TGF-*β*1 and ECM remodeling. Each cell was represented as a point in the system with an integrated system of ODEs that simulated changes in TGF-*β*1 -induced intracellular signaling pathways. Changes in cell state were determined by changes in the EMT marker N-Cadherin. This model provides a predictive tool to simulate the effects of exogenous TGF-*β*1 and fibrotic ECM assembly on EMT progression of a multi-cell population. The model captured localized TGF-*β*1 production by cells, as well as tethering in areas of low diffusion coefficient values. Previous studies have shown that increased FN fibrillogenesis reveals cryptic binding sites that bind growth factors including TGF-*β*1, creating reservoirs of growth factors in the microenvironment [23, 48]. We show clear colocalization of high TGF-*β*1 concentration at areas of low local diffusion coefficient values, representative of increased fibril formation that contributes to this fibrotic phenotype.

Results indicate that increasing fibrotic ECM, modeled as a decrease in local TGF-*β*1 diffusion, drives cells towards a partial EMT state in both the presence and absence of exogenous TGF-*β*1. Results indicate that cells do not lose their spheroid morphology with exposure to exogenous TGF-*β*1, but instead experience a dilation of the spheroid, in which there is a measurable increase in lumen size. Both of these findings are in agreement with pre-clinical and clinical research. A main characteristic of tubulointerstitial fibrosis in the kidney is tubule dilation, where epithelial cells that comprise tubules and collecting ducts in the nephron depolarize, lose cell-cell junctions, and increase in tubule diameter [53]. Interestingly, the dilation of these epithelial structures is driven in large part by the transcription factor Snail1 [54–56], which is consistent with results from our model in which Snail1 is significantly upregulated in response to fibrotic ECM.

There is significant evidence in several disease states, including kidney fibrosis, cardiovascular disease, and cancer, that indicates a predominance of epithelial cells in a partial EMT state. In kidney fibrosis, there is active debate regarding the role of EMT; the discussion has focused on whether or not tubular epithelial cells differentiate into myofibroblasts, which are considered the main producers of fibrotic ECM [14, 57, 58]. Studies have indicated that *in vivo*, tubular epithelial cells remain in a partial EMT state and do not contribute to the myofibroblast population during fibrosis [59]. Instead, these epithelial cells perform an indirect role in promoting fibrogenesis, through secretion of profibrotic growth factors and depolarization of epithelial structures that progresses to kidney failure. Similarly, this can also be seen in the vasculature, where endothelial cells undergo a similar process known as EndoMT, where endothelial cells progressively lose cell-cell junctions that can lead to vascular rarefaction known to contribute to fibrogenesis and overall organ failure [60]. In the context of metastasis, studies have indicated that circulating tumors are composed of a heterogeneous population of cells in a partial EMT state, allowing for invasion between basement membrane and vasculature and promoting collective cell migration. Further discussion has suggested that cells in a partial EMT state show increased survival rates and increased resistance to cancer therapeutics [8, 61]. Our findings that a fibrotic ECM drives epithelial cells to a partial EMT state, regardless of exogenous TGF-*β*1 concentration, are consistent with these findings and suggest a mechanism through which fibrotic ECM could facilitate these disease states.

There are limitations in using an agent-based model, such as a lack of distinct representation of changes in cell size and cell-cell junction integrity, which are critical in epithelial tissue organization. Although the use of a Cellular Potts model to study cell-cell interactions has been the gold standard due to its ability to model changes in cell shape, size, and junctional forces [19, 35], adaption into 3D system requires high computational processing and memory as well as increased complexity of the model itself. As such, we chose an agent-based model to represent cells in a 3D system, introducing probability thresholds to limit cell movement. This also allows us to closely study TGF-*β*1 diffusion and interactions between TGF-*β*1 and the surrounding ECM.

Future studies will further explore cell-matrix interactions in EMT, with particular focus on 3 areas: i) incorporation of multiple cell types; ii) incorporation of multiple spheroids, and iii) simulation of therapeutic interventions for preventing EMT within spheroids. The current model only simulates epithelial cells within the spheroid structure; however, multiple cell types in the surrounding interstitial space contribute to ECM remodeling and TGF-*β*1 signaling [62–64]. Future iterations will incorporate these cells into the extra-spheroid interstitial space, which will allow for dynamic changes to fibrotic ECM as opposed to the constant initial conditions used here. Current simulations also utilized single spheroids; however, *in vivo*, tubules in the kidney and ducts in mammary tissue are closely packed and may have synergistic effects on neighboring structures. In addition, many *in vitro* spheroid assays involve multiple spheroids embedded into a single hydrogel [65]. Future studies will incorporate multiple spheroids into the computational space in order to identify potential synergistic and spatiotemporal effects of neighboring spheroids. Additionally, the current model is initialized with all cells in an epithelial state; future work will integrate a heterogeneous population of initial EMT states to improve physiological relevance. In terms of therapeutic interventions, we can use this model to study treatment methods associated with ECM production and degradation based on D_TGF-*β*1_ value. Targeting the ECM to treat fibrotic diseases has shown promising results: pUR4 is a protein fragment that inhibits fibronectin fibril assembly, and numerous *in vivo* studies have shown that it is able to ameliorate fibrosis in cardiac disease, liver, kidney, and in myopic fibrosis [15,25].

## Conclusions

Difficulties in treating fibrotic diseases stem from the complex interactions between cells and the surrounding ECM. The development of a 3D agent-based model allows us to simulate a 3D epithelial cell tissue structure typically used *in vitro* to study fibrosis and other epithelial-associated diseases. The integration of ECM remodeling by changing the TGF-*β*1 diffusion coefficient is able to reproduce cell behaviors seen *in vitro* without direct ECM quantification or kinetics. Both spatial and temporal dynamics of TGF-*β*1 signaling are important considerations for EMT progression in chronic disease states like renal fibrosis and lung fibrosis, further highlighting the important role of cell-ECM interactions as key mechanisms of disease progression. We demonstrate that the use of this agent-based model simulates the progression of EMT in 3D epithelial spheroids, where increasing exogenous TGF-*β*1 concentrations drove increased spheroid dilation and increased mesenchymal markers. Incorporation of a fibrotic ECM by locally decreasing the TGF-*β*1 diffusion predicted alterations to EMT state, indicating that fibrotic ECM is a major factor that drives cells in the epithelial spheroid towards a partial state, regardless of exogenous TGF-*β*1 concentration.

## Acknowledgments

This research was supported through the IGNITE KUH Training Fellowship. We would also like to thank Samantha Ho for help with performing simulations and developing future applications of the model.

## Supporting information

**Table S1. Intracellular TGF-*β*1 signaling equations** The system of ordinary differential equations that govern TGF-*β*1 signaling and subsequent EMT characterization for each individual cell, derived from Tian et al (2013).

**Table S2. Intracellular TGF-*β*1 signaling reaction rates.** Reaction rates and other values used to define intracellular TGF-*β*1 signaling dynamics.

**Table S3. Intracellular TGF-*β*1 signaling parameters. Initial values of time-dependent species modeled.**

**Table S4. Model parameters describing the extracellular matrix environment and cell characteristics.** D_TGF-*β*1_ represents the TGF-*β*1 diffusion coefficient, where the bolded value represents the normal ECM. Cell size defines the size of the system. Cell migration and proliferation depend on the probability (P) of events occurring, set to change based on the EMT state [epithelial, partial, mesenchymal]. Probability thresholds were adapted from Hirway et al (2021).

**Movie S1. Progression of EMT in spheroids at 0 TGF-***β***1, 38 D_TGF-_***_β_***_1_ values.** A video output that shows 24 hour time intervals of the spheroid up to 10 days/ 260 hours. Baseline parameters were used and spheroids were initialized in healthy ECM (D_TGF-*β*1_ = 38 pixels^2^/hr) and with no exogeneous TGF-*β*1. EMT states are denoted as epithelial (green), partial (yellow), and mesenchymal (red).

**Movie S2. Progression of EMT in spheroids at 2 µM TGF-***β***1, D_TGF-_***_β_***_1_ = 38 pixels^2^/hr.** A video output that shows 24 hour time intervals of the spheroid up to 10 days/ 260 hours. Baseline parameters were used and spheroids were initialized in healthy ECM (D_TGF-*β*1_ = 38 pixels^2^/hr) and in the presence of 2 µM TGF-*β*1. This video shows changes in cell position and EMT state, as well as changes in TGF-*β*1 gradient concentration and local D_TGF-*β*1_ values. EMT states are denoted as epithelial (green), partial (yellow), and mesenchymal (red).

**Movie S3. Progression of EMT in spheroids at 4 µM TGF-***β***1, D_TGF-_***_β_***_1_ = 38 pixels^2^/hr.** A video output that shows 24 hour time intervals of the spheroid up to 10 days/ 260 hours. Baseline parameters were used and spheroids were initialized in healthy ECM (D_TGF-*β*1_ = 38 pixels^2^/hr) and in the presence of 4 µM TGF-*β*1. This video shows changes in cell position and EMT state, as well as changes in TGF-*β*1 gradient concentration and local Dtgfb values. EMT states are denoted as epithelial (green), partial (yellow), and mesenchymal (red).

**Movie S4. Progression of EMT in spheroids at 0 µM TGF-***β***1, D_TGF-_***_β_***_1_ = 0 pixels^2^/hr.** Output of the model without exogenous TGF-*β*1, and spheroid initialized in a fibrotic ECM environment (D_TGF-_*_β_*_1_ = 0 pixels^2^/hr). Video shows 24 hour time intervals up to 10 days/ 260 hours. EMT states are denoted as epithelial (green), partial (yellow), and mesenchymal (red).

**Movie S5. Progression of EMT in spheroids at 4 µM TGF-*β*1, D_TGF-*β*1_ = 0 pixels^2^/hr.** Output of the model with an added 4 µM concentration of exogenous TGF-*β*1, and spheroid initialized in a fibrotic ECM environment(D_TGF-*β*1_ = 0 pixels^2^/hr). Video shows 24 hour time intervals up to 10 days/ 260 hours. EMT states are denoted as epithelial (green), partial (yellow), and mesenchymal (red).

**Figure S1. Heat map of RMSD error values when comparing discrete combinations of K1 and K3 at different exogenous TGF-*β*1 concentrations.** Average RMSD was found by taking the average of RMSD outputs at the 4 different exogenous TGF-*β*1 concentrations.

**Figure S2. Heat map of average RMSD error values when comparing discrete combinations of K1 and K3.** Average RMSD was found by taking the average of RMSD outputs at the 4 different exogenous TGF-*β*1 concentrations.

**Figure S3. Changes in straight tube morphology in response to different TGF-*β*1 concentrations.** Outputs comparing EMT progression and changes in the straight tubular morphology at different exogenous TGF-*β*1 concentrations. A. Representative images of tubular structures, TGF-*β*1 gradient, and D_TGF-*β*1_ gradients. B. Cell morphology (total cell count, cross-sectional area). C. Changes in EMT Progression (EMT population ratio, EMT markers (E-Cadherin, Snail, N-Cadherin)), D. Changes in cell proliferation and organization over time. E. Changes in EMT marker concentration over time. N = 20 simulations per condition. Statistics were run using a 1-way ANOVA and post hoc Tukey multiple comparison test (*^∗^p ≤* 0.033, *^∗∗^p ≤* 0.002, *^∗∗∗^p ≤* 0.001).

**Figure S4. Changes in curved tube morphology in response to different TGF-*β*1 concentrations** Outputs comparing EMT progression and changes in the curved tubular morphology at different exogenous TGF-*β*1 concentrations. A. Representative images of tubular structures, TGF-*β*1 gradient, and D_TGF-*β*1_ gradients. B. Cell morphology (total cell count, cross-sectional area). C. Changes in EMT Progression (EMT population ratio, EMT markers (E-Cadherin, Snail, N-Cadherin)), D. Changes in cell proliferation and organization over time. E. Changes in EMT marker concentration over time. N = 20 simulations per condition. N = 20 simulations per condition. Statistics were run using a 1-way ANOVA and post hoc Tukey multiple comparison test (*^∗^p ≤* 0.033, *^∗∗^p ≤* 0.002, *^∗∗∗^p ≤* 0.001)

**Figure S5. MATLAB App Designer Code Organization.** To improve future application of our model in other research, we developed a GUI using the MATLAB App Designer. An example of the user interface. Users are able to change parameters of cell morphology, TGF-*β*1 signaling, ECM composition, and total time the model is run. We further allow several methods of data collection where users are able to save outputs of multiple runs in excel files, or save graphical results as image files (.jpg or .tif) or video format (.gif).

